# MLcps: Machine Learning Cumulative Performance Score for classification problems

**DOI:** 10.1101/2022.12.01.518728

**Authors:** Akshay Akshay, Masoud Abedi, Navid Shekarchizadeh, Fiona C. Burkhard, Mitali Katoch, Alex Bigger-Allen, Rosalyn M. Adam, Katia Monastyrskaya, Ali Hashemi Gheinani

**Affiliations:** Functional Urology Research Group, Department for BioMedical Research DBMR, University of Bern, Switzerland; Graduate School for Cellular and Biomedical Sciences, University of Bern, Switzerland; Department of Medical Data Science, Leipzig University Medical Centre, 04107 Leipzig, Germany; Center for Scalable Data Analytics and Artificial Intelligence (ScaDS.AI) Dresden/Leipzig, 04105 Leipzig, Germany; Department of Urology, Inselspital University Hospital, 3010 Bern, Switzerland; Institute of Neuropathology, Universitätsklinikum Erlangen, Friedrich-Alexander-Universität Erlangen-Nürnberg (FAU), Erlangen, Germany; Biological & Biomedical Sciences Program, Division of Medical Sciences, Harvard Medical School, Boston, MA; Urological Diseases Research Center, Boston Children’s Hospital, MA, USA; Harvard Medical School, Boston, Department of Surgery MA, USA; Broad Institute of MIT and Harvard, Cambridge, MA, USA

**Author notes:** Corresponding author: Ali Hashemi Gheinani, Urological Diseases Research Center, Boston Children’s Hospital, Harvard Medical School and Broad Institute of MIT and Harvard, Cambridge, MA, USA.

## Abstract

**Motivation:** A performance metric is a tool to measure the correctness of a trained Machine Learning (ML) model. Numerous performance metrics have been developed for classification problems making it overwhelming to select the appropriate one since each of them represents a particular aspect of the model. Furthermore, selection of a performance metric becomes harder for problems with imbalanced and/or small datasets. Therefore, in clinical studies where datasets are frequently imbalanced and, in situations when the prevalence of a disease is low or the collection of patient samples is difficult, deciding on a suitable metric for performance evaluation of an ML model becomes quite challenging. The most common approach to address this problem is measuring multiple metrics and compare them to identify the best-performing ML model. However, comparison of multiple metrics is laborious and prone to user preference bias. Furthermore, evaluation metrics are also required by ML model optimization techniques such as hyperparameter tuning, where we train many models, each with different parameters, and compare their performances to identify the best-performing parameters. In such situations, it becomes almost impossible to assess different models by comparing multiple metrics.

**Results:** Here, we propose a new metric called Machine Learning Cumulative Performance Score (MLcps) as a Python package for classification problems. MLcps combines multiple pre-computed performance metrics into one metric that conserves the essence of all pre-computed metrics for a particular model. We tested MLcps on 4 different publicly available biological datasets and the results reveal that it provides a comprehensive picture of overall model robustness.

**Availability:** MLcps is available at https://pypi.org/project/MLcps/ and cases of use are available at https://mybinder.org/v2/gh/FunctionalUrology/MLcps.git/main.

**Supplementary information:** Supplementary data are available at Bioinformatics online.

## Introduction

Performance metrics play a crucial role in both evaluating model performance and model optimization [1]. Different performance metrics are widely available to evaluate and compare the performance of ML classification models. However, previous research has shown that a model that performs well based on a particular metric may not perform well on another metric [2-4]. Supervised learning algorithms (such as classification or regression problems), and unsupervised learning algorithms (such as clustering), have different evaluation metrics according to their outputs. In addition, most performance metrics are sensitive to the composition of the dataset under study [5].

In practice, it is not enough to measure only one metric to assess the overall performance of a model because most of the metrics give importance to a particular aspect of the model’s performance. In fact, in many practical applications, there is an unavoidable trade-off between these performance measures. Therefore, ranking a classifier “superior” in terms of a particular metric can result in a relatively “inferior” classifier in terms of another. For example, the “recall” metric can summarize how well a model can predict the positive class but gives no information about the negative class. Therefore, it becomes essential to measure and compare different performance metrics for each model to identify the best-performing model.

Although it is reasonable to expect that the best-performing model would have the highest score for all the measured metrics, this is an optimistic assumption and unlikely to be true in practice. Comparison of different metric scores for a large number of models to identify the best-performing model is labour-intensive, and this process is prone to user preference bias [6]. Apart from that, during the model optimization step, the goal is to identify the best-performing parameter for a model. For this purpose, a very common approach in the ML field is to train a large number of models with different parameters. In such a situation, finding the best-performing model by comparing different metrics becomes very complex.

To address this problem, we propose a new metric called Machine Learning Cumulative Performance Score (MLcps) for classification problems that combines multiple pre-computed performance metrics into one metric while preserving the essence of all pre-computed metrics for a particular model. MLcps is available as a Python package and allows the straightforward comparison of trained ML models to rank the model’s performance. The input of MLcps package is a table containing the names of classification ML algorithms and their relevant individual performance scores. The output of the package is a table and the relevant bar chart representing different ML algorithms, ranked based on a cumulative performance score, calculated by MLcps.

## MLcps Methodology

The input for the MLcps package is a table in which columns contain metrics (such as F1, Accuracy, and Recall) and the rows contain ML methods of choice (such as K-Nearest Neighbors (KNN) and Support-Vector Machine (SVM)). This table should be generated after performing feature extraction on raw data (Figure 1.A), training ML algorithms (Figure 1.B), and performance evaluation (Figure 1.C). To calculate MLcps, first, we draw all pre-calculated performance metrics on a two-dimensional polar coordinates system (Figure 1.D), in which each point on a plane represents an individual metric and is determined by the distance from a reference point that is a pole and an angle from a reference direction. This is similar to a radar graph, which is a circular graphing method and has a series of rays projecting from a reference point, with each ray representing a different metric label (Figure S1-S2).

**Figure 1.**
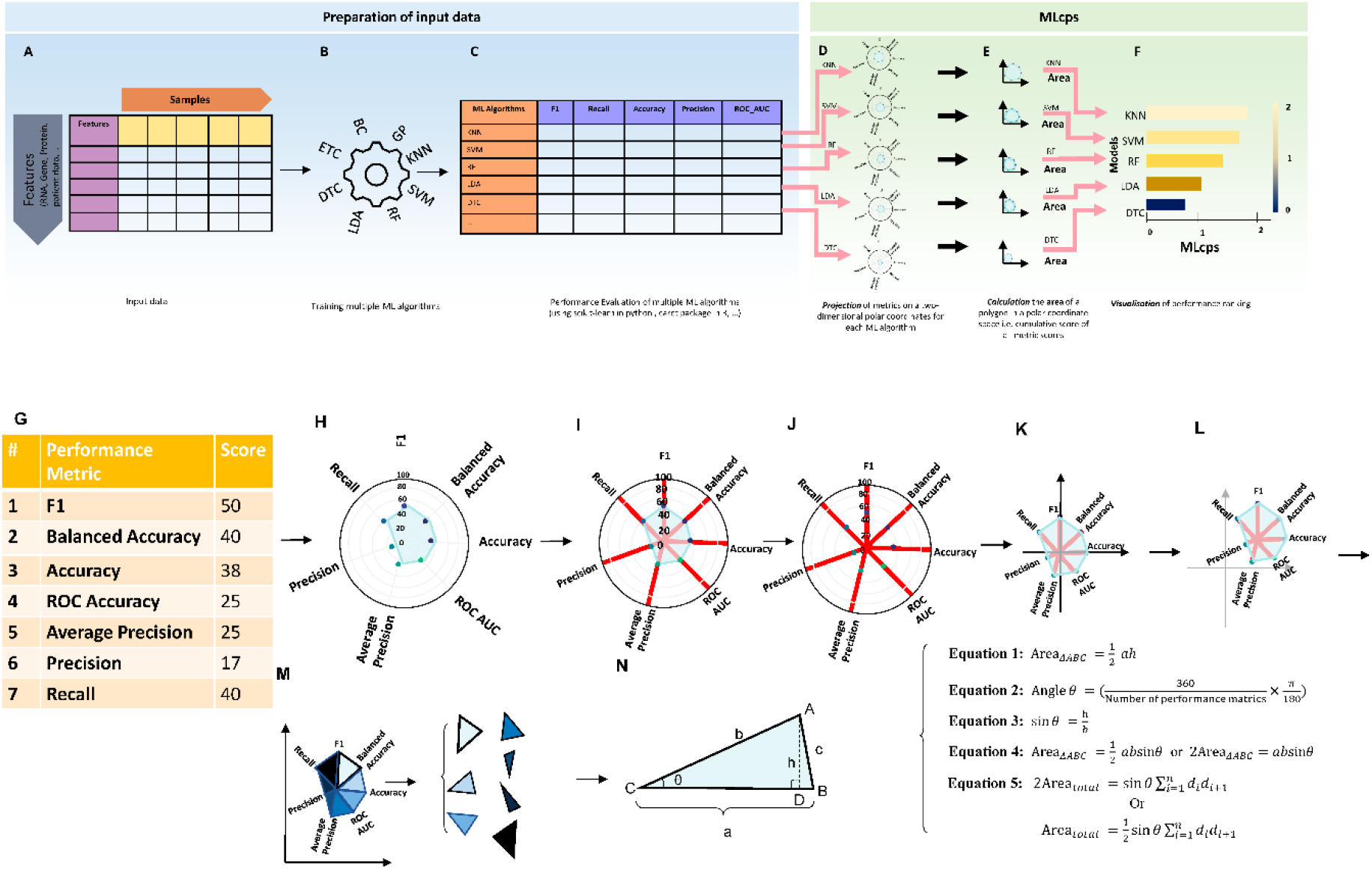
Schematics of a full analysis process for MLcps Python package. Before using the MLcps Python package, one needs to prepare the raw data (A). This input table can be RNA sequencing, proteomics, patients’ profile, molecular data, etc (normally this data is in txt or csv format). Next step is to perform multiple ML algorithms (B). Performing this step can be done by any package or programming language of choice. The next step is to evaluate the performance of the ML algorithms. We recommend the use of multiple metrics such as F1, Recall, etc (C). The performance metric scores then need to be arranged in a tabular format as depicted in (C). This table will be used as an input for the MLcps package. From here on the MLcps will process the data. MLcps involves three steps: projection, calculation, and visualization (PCV). To calculate the cumulative score of each ML algorithm in the input data, MLcps first projects the performance metric onto the two-dimensional polar coordinates system (D). Next, the projected polygon’s area is calculated (E). Finally, the user can visualize this MLcps to rank the performance of given ML algorithms (F). The lower panel (G-N) visualises the procedure to calculate the surface area as cumulative score in detail. The names of the algorithms are just mentioned as example and other algorithm can be used too. ETC: Extra Trees Classifier, BC: Bagging Classifier, GP: Gaussian Process Classifier, KNN: K-Nearest Neighbors, SVM: Support Vector Machine, RF: Random Forest Classifier, LDA: Linear Discriminant Analysis, DTC: Decision Tree Classifier.

In the polar plane, the values of the metrics are encoded into the lengths of the rays. To generate a cumulative score, we compute the area of each polar plane (Figure 1.E). We assume that each plane is divided into multiple triangles and the sum of the individual triangles is the area of the whole plane. In addition, the area of the created surface on the polar plane is proportional to the cumulative magnitude of all metrics being analysed. Finally, the cumulative performance score (cps) is visualized as a bar chart (Figure 1.F).

### Area calculation of a two-dimensional polar surface

To calculate the cumulative performance score, we need to know the total area of a polar plane generated by multiple performance scores. The surface is then divided into multiple triangles and the sum of the area of individual triangles is the area of the whole plane (cumulative performance score). As the general formula for the area of a triangle is stated in Equation 1: the area of a triangle equals ½ the length of one side times the height drawn to that side (Figure 1.G-N).

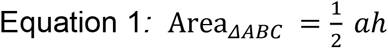

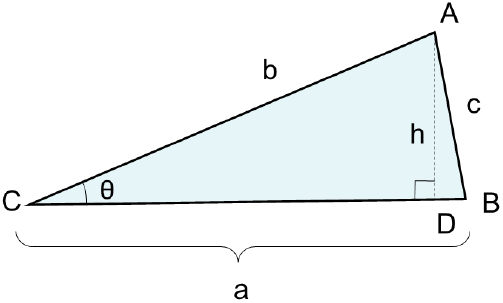

where *a* represents the side (base)

and *h* represents the height drawn to that side.

To use this formula, the *h* is needed, which cannot be controlled in radar graph, whereas the angle of all triangles can be controlled by dividing 360 by the number of used performance metrics:

The angle between each two-performance metric on a radar graph is:

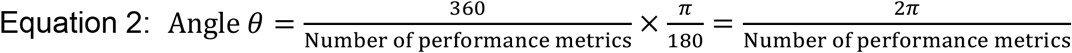

Therefore, using trigonometry we will rewrite the area formula and express the height as follows:

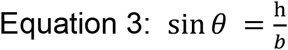

Therefore, the height h of the triangle can be expressed as *b sin θ*. Now substituting for the height h into the general formula for the area of a triangle gives:

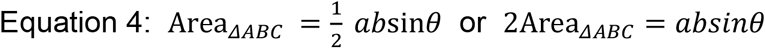

where *a* and *b* can be any two sides and *θθ* is the included angle. The parameters *a* and *b* are indeed the actual values for each performance metric.

Thus, the sum of areas of all triangles forming the polar coordinate is computed based on the following formula:

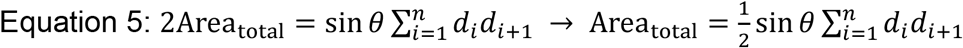

Where:d_i_ = length of the ith ray (the value of ith metric score) (Figure 1L) n = number of triangles point of collapse (Figure 1M)

### Weighted MLcps

In particular cases, certain metrics are more significant than others. For example, if the dataset is imbalanced, a high F1 score might be preferred over higher accuracy [7]. A user can provide weight variables for metrics of interest while calculating MLcps in such a scenario. A weight variable provides a value (the weight) for each pre-computed metric and the given metrics score will be updated using these weights in the following way:

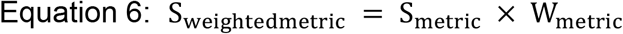

Where:

S_weightedmetric_ = Weighted metric Score

S_metric_ = Raw metric Score

W_metric_ = Weight

The given weight for a metric should always be greater than or equal to zero. Zero weight implies that the user wants to exclude it from the MLcps. Metrics with relatively large weights contribute more to the MLcps than metrics that have smaller weights. An unweighted MLcps is equal to a weighted analysis in which all weights are 1.

## Results and discussion

### Testing MLcps on 4 different real-world dataset examples

We have used four different datasets to test MLcps (*Table 1*). The first dataset is an mRNA dataset (n=136) from a study of chronic lymphocytic leukemia (CLL) that measured transcriptome profiles in blood cancer patients [8]. Here we aimed to train a model to classify male and female patients based on their transcriptomic profiles. For that, we considered the top 5,000 most variable mRNAs after the exclusion of genes from the Y chromosome.

**Table 1.**
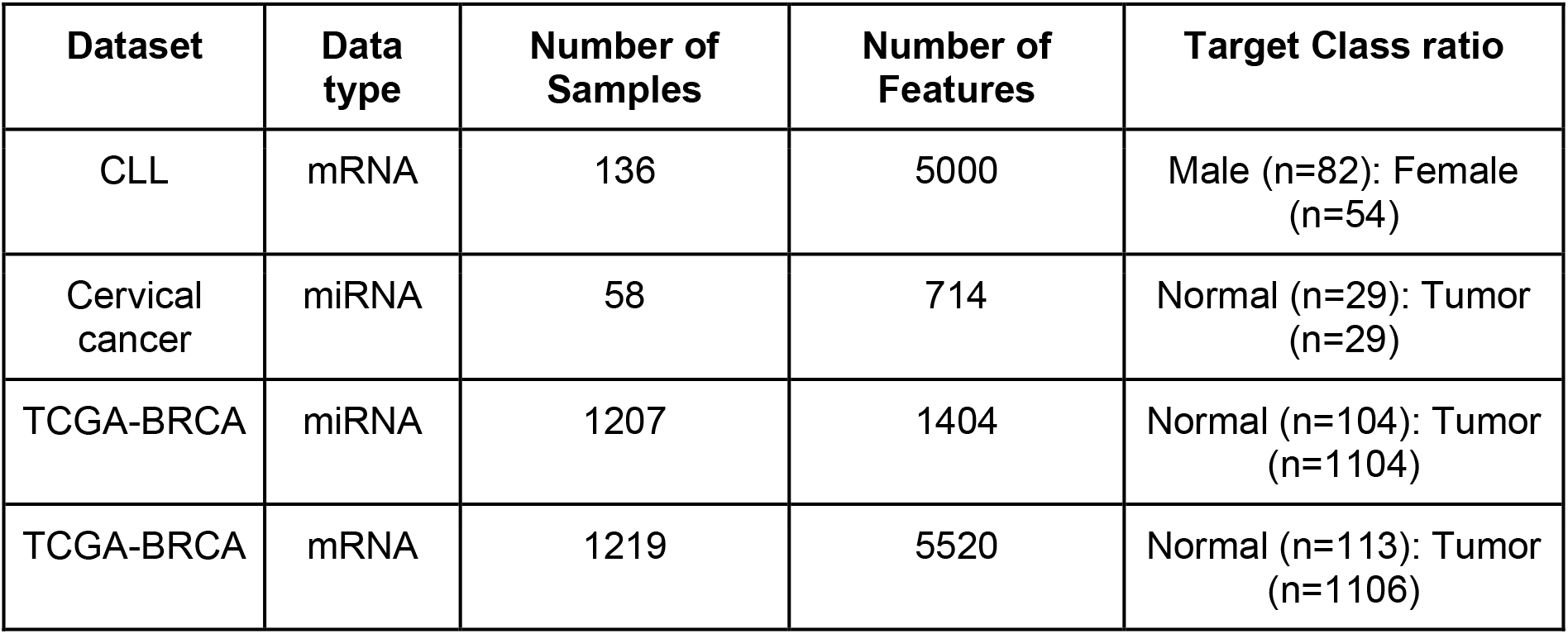
Example datasets used in this study

| Dataset | Data type | Number of Samples | Number of Features | Target Class ratio |
| --- | --- | --- | --- | --- |
| CLL | mRNA | 136 | 5000 | Male (n=82): Female (n=54) |
| Cervical cancer | miRNA | 58 | 714 | Normal (n=29): Tumor (n=29) |
| TCGA-BRCA | miRNA | 1207 | 1404 | Normal (n=104): Tumor (n=1104) |
| TCGA-BRCA | mRNA | 1219 | 5520 | Normal (n=113): Tumor (n=1106) |

The second set of data is taken from a cervical cancer study that measured the expression levels of 714 miRNAs in human samples (n=58) [9]. The third and fourth datasets are retrieved mRNA (n=1219) and miRNA (n=1207) sequencing of Breast Invasive Carcinoma (BRCA) from The Cancer Genome Atlas (TCGA) using the TCGAbiolinks package [10] in R. For the BRCA mRNA dataset, we only considered differentially expressed genes from edgeR (FDR <= 0.001 and logFC > ±2) [11]. We aimed to train a model that distinguishes between normal and tumor samples for cervical cancer and TCGA-BRCA datasets. Two of these datasets are comparatively small datasets (CLL and a cervical cancer study) and the remaining two are imbalanced datasets (*Table 1*). For all datasets, we trained and evaluated 8 different models (*Table S1*) using an in-house developed ML pipeline to find out the best-performing models (Figure 2).

**Table S1.**
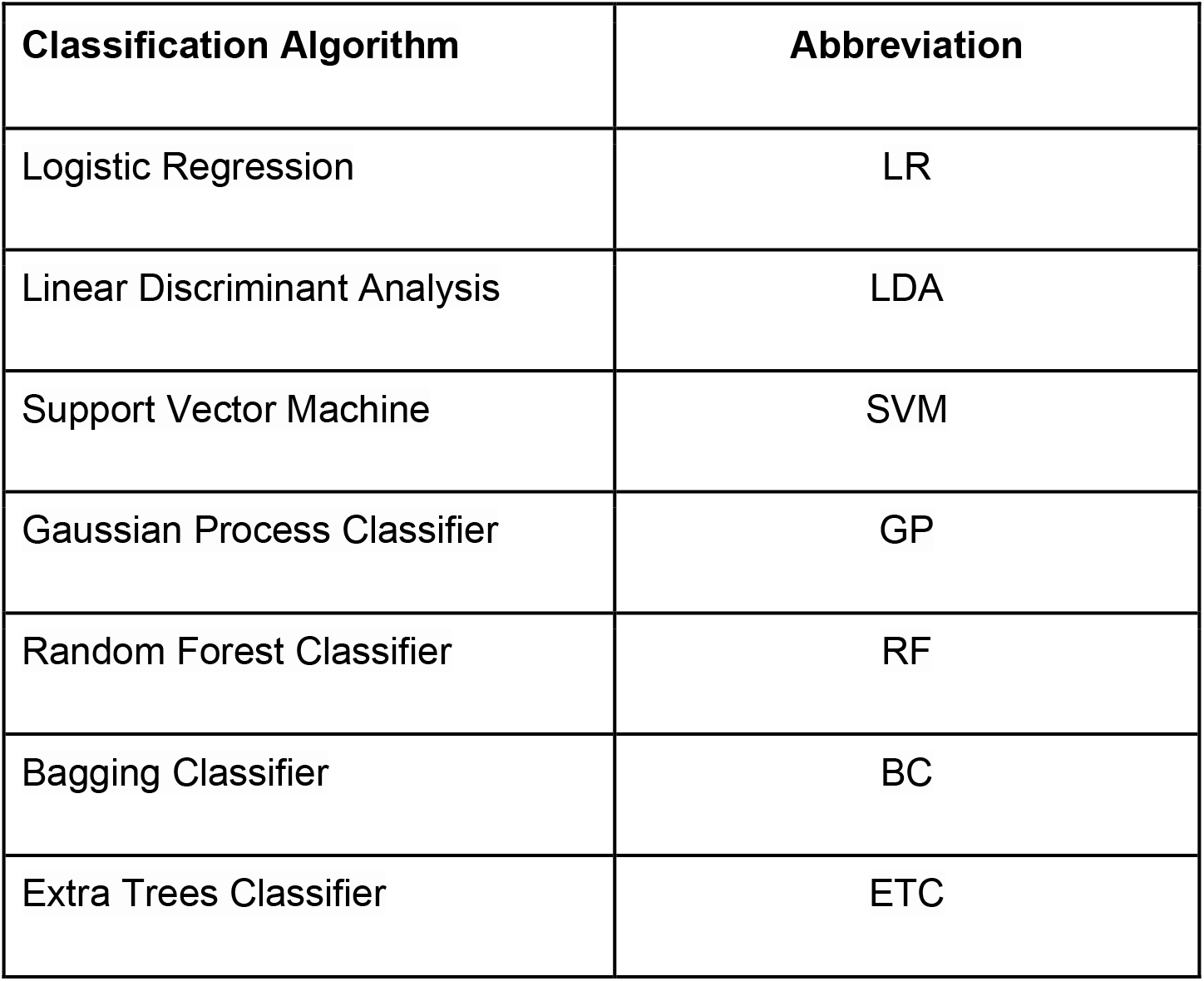
ML algorithms used in this study

**Figure 2.**
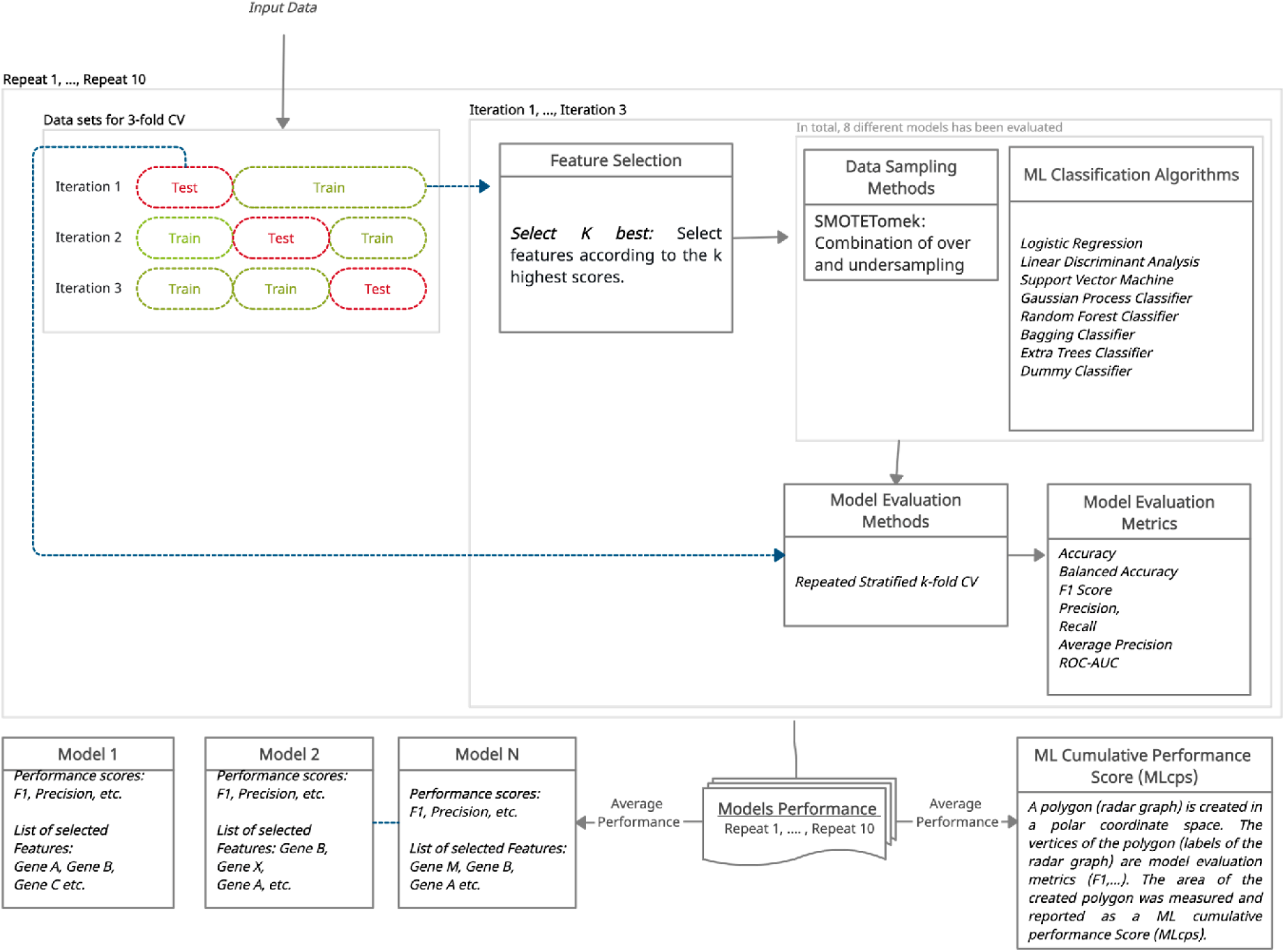
Flowchart describing ML Pipeline: The Used ML pipeline first splits the datasets into k (3) equal size different bins in a stratified manner, where k-1 bins will be used as training datasets and the remaining bin as a test dataset. Next, it uses the univariate feature selection method for feature selection, SMOTETomek method for data resampling, and eight ML algorithms. Then, it evaluates the model performance on the test dataset using the k-fold and nested CV (k=3) method and calculates seven different performance metrics. As part of k-fold CV methods, it repeats the last two steps for each unique bin. It repeats the complete process ten times and takes average performance as a final model performance. In the end, it provides a list of the intersection of selected features from top 10 best performing models (based on F1 score) as a final list of selected features.

### Testing MLcps robustness based on the standard deviation of performance metrics

For all datasets, except the TCGA mRNA BRCA dataset, the model with the highest MLcps has the lowest Standard Deviation (SD) in the performance metrics score (*Figure 3 A-D, Figure 4 A-F*). The Dummy model for the CLL dataset has low SD compared to the GP model (*Figure 3A*) and the LDA model for the cervical cancer dataset has low SD compared to the ETC, SVM, and RF models (*Figure 3B*). However, the MLcps of GP model based on the CLL dataset is higher compared to the corresponding MLcps of Dummy model, and the MLcps of ETC, SVM, and RF models for the cervical cancer dataset is higher compared to MLcps of LDA model (*Figure 4A & 4B*). It confirms that, besides SD, MLcps also considers the overall magnitude of performance metric score. MLcps tends to nominate a model as the best one when it has the lowest SD in metrics score but an overall high metrics score compared to other models. In *Figure 3D*, LR would be chosen as the best-performing model when relying entirely on SD. However, LR is not even in the top 3 if we look at its performance on the test dataset (*Figure 4F*). On the other hand, if we sort the performance of models based on MLcps, the order is almost identical on training as well as test dataset (*Figure 4C & 4D, Figure 4E & 4F*).

**Figure 3.**
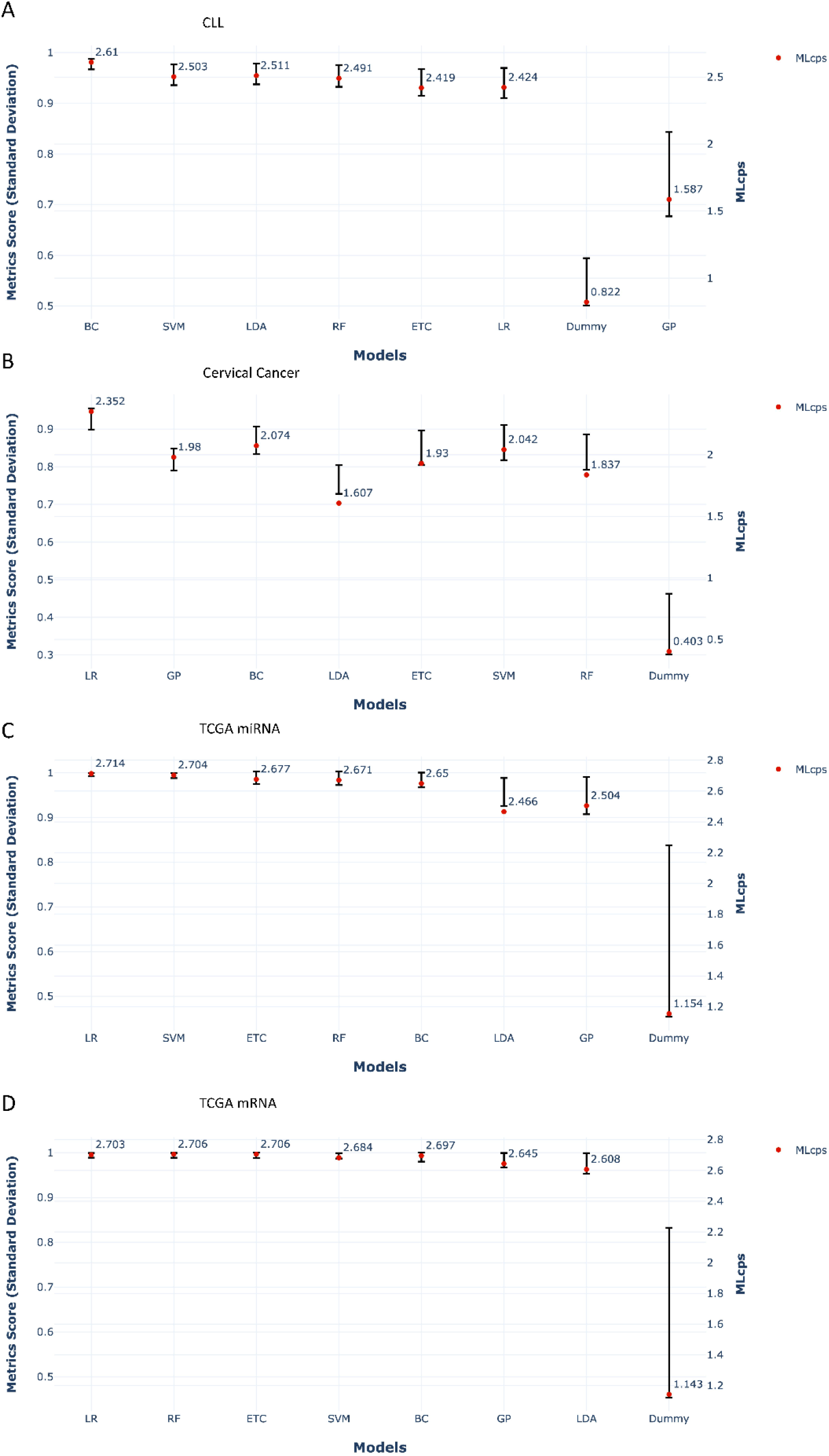
Standard deviation (SD) of performance metrics score of ML algorithms trained on different example datasets. Plots are representing A) CLL dataset, B) Cervical cancer dataset, C) TCGA miRNA dataset, and D) TCGA mRNA dataset. Each bar in a plot represents SD of performance metrics score (left y-axis), and red dot represents MLcps (right y-axis).

**Figure 4.**
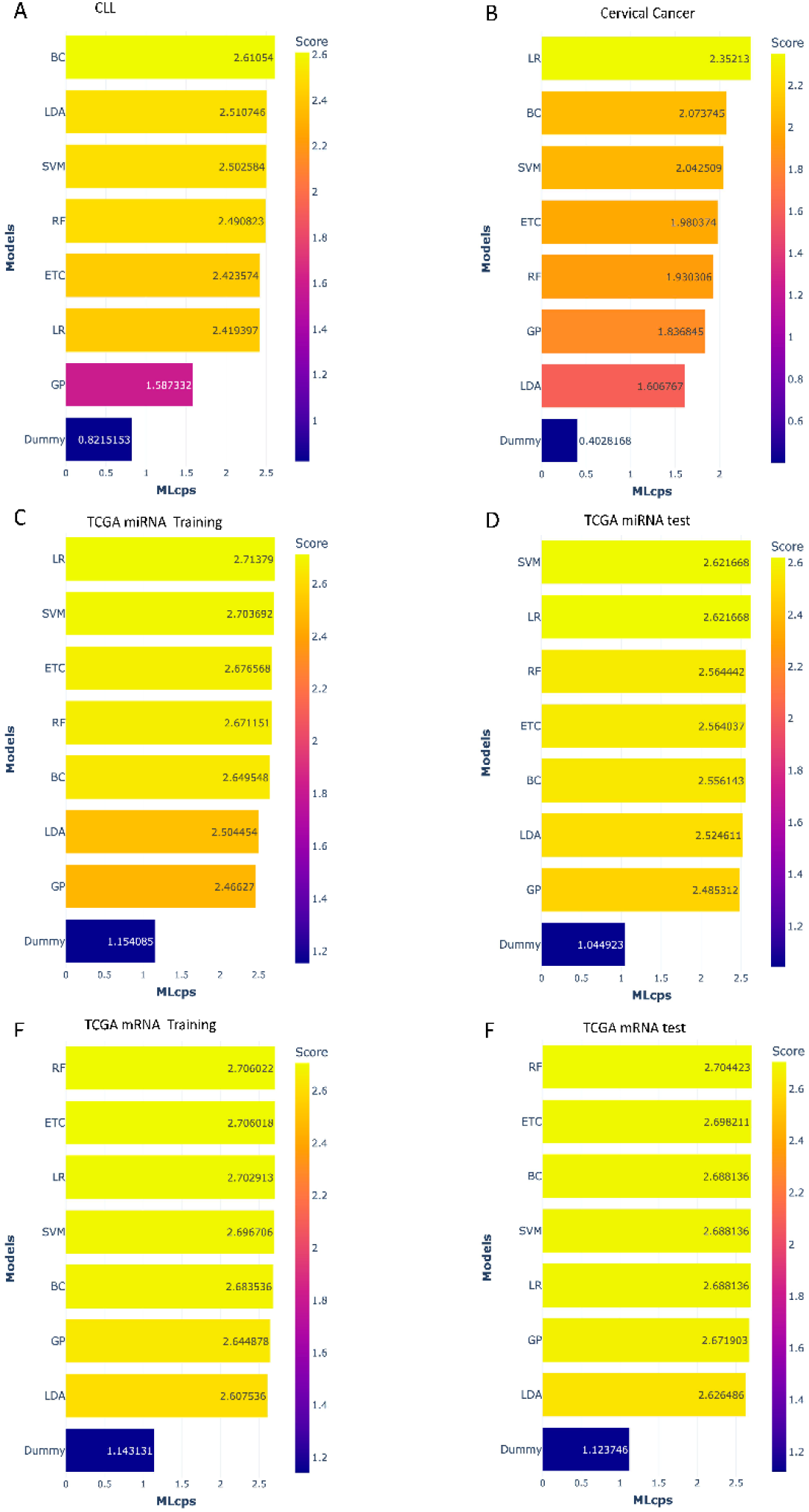
Bar charts visualizing the MLcps for different example datasets. Plots are representing A) CLL dataset, B) Cervical cancer dataset, C) TCGA miRNA training dataset, D) TCGA miRNA test dataset, E) TCGA mRNA training dataset, and F) TCGA mRNA test dataset. The color and the length of the bars represent the MLcps. The darker the color, the lower the score

### Comparing the ML ranking in training and testing datasets

The TCGA-BRCA datasets are larger and thus allow us to create an independent testing set (30% of the datasets). Here the best-performing model selected based on MLcps also shows the best performance when applied to an independent test set (*Figure 4. C-F*).

### Visualization of the importance of using multiple performance metrics

In Figures, S1B & C, and Figure S2B & C, the precision and average precision performance metrics showed high scores (>90%) even in the dummy model for TCGA mRNA and miRNA datasets. We would have selected the wrong model as the best-performing one if we had used only precision and average precision metrics to evaluate the model performance for TCGA datasets. It demonstrates the importance of using multiple performance metrics to assess ML model performance accurately. This phenomenon is not observed in the CLL and cervical cancer datasets (*Figure S1A, Figure S2A*).

**Figure S1.**
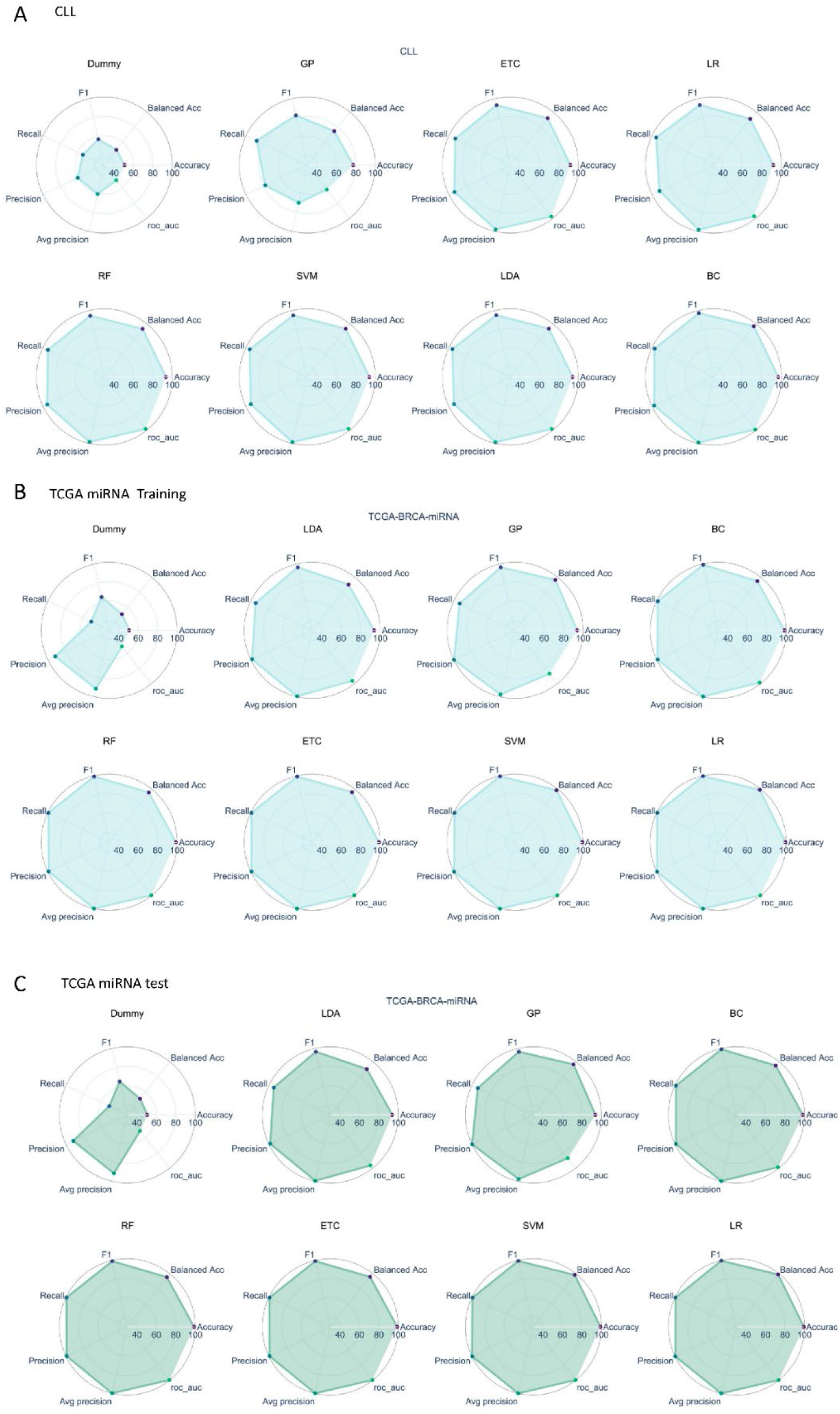
Projection of metrics scores on two-dimensional (2D) polar coordinates for each ML algorithm trained on different example datasets. Plots are representing A) CLL dataset, B) TCGA miRNA Training dataset, C) TCGA miRNA test dataset.

**Figure S2.**
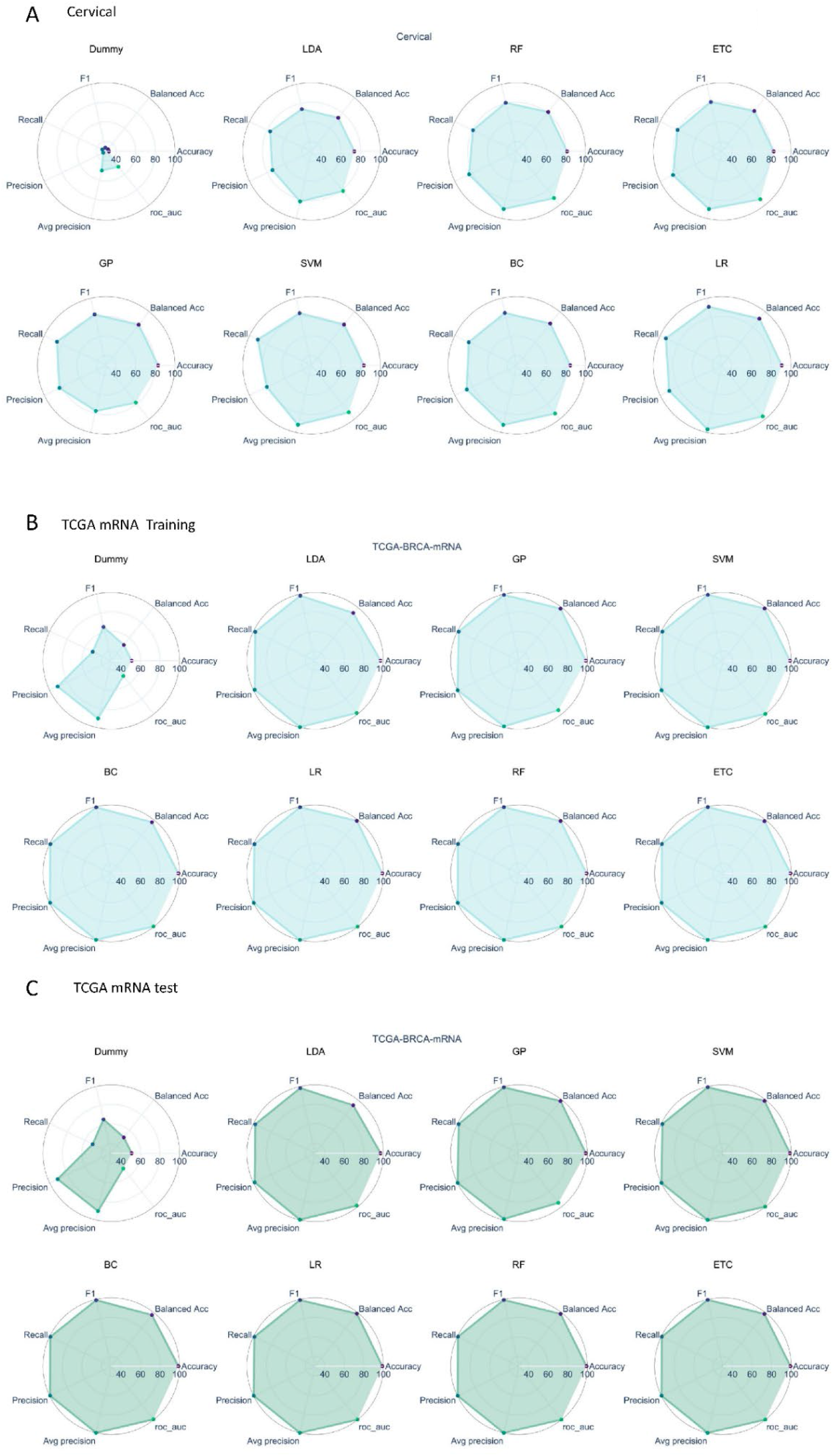
Projection of metrics score on a two-dimensional (2D) polar coordinate for each ML algorithm trained on different example datasets. Plots are representing A) Cervical cancer dataset, B) TCGA mRNA Training dataset, C) TCGA mRNA test dataset.

### Selection criteria for best-preforming Model

Considering that it is highly unlikely to find a model with the highest score across all measured metrics, the use of multiple metrics makes it more difficult to find the best-performing model. Alternatively, one could define a set of criteria, and a model that best meets those criteria would be considered as the best-performing model. In line with the latter approach, we have defined the following criteria for selecting the best-performing model:

- **Standard Deviation (SD) in performance metric score:** Since each of the performance metrics represent a particular aspect of the model’s performance, the most robust overall model will have similar scores for most of the metrics. Therefore, compared to other models, the best-performing model should have a small SD in the performance metrics score (*Figure 3 A-D*).
- **Overall magnitude of performance metric score:** There is a possibility that a model with poor performance metrics could have a smaller SD compared to other models. Hence, one cannot rely solely on the SD criterion and should also consider the overall magnitude of the metric score (*Figure 3 A-D*).

### Usage

Note: If you want to use MLcps without installing it on your local machine, please click here https://mybinder.org/v2/gh/FunctionalUrology/MLcps.git/main. It will launch a Jupyterlab server (all the requirements for MLcps are pre-installed) where you can run the already available example Jupyter notebook for MLcps analysis. It may take a while to launch! You can also upload your data or notebook to perform the analysis.

### Prerequisites

- Python >=3.8
- R >=4.0
- radarchart, tibble, and dplyr R packages. MLcps can install all these packages at first import if unavailable, but we highly recommend installing them before using MLcps. The user could run the following R code in the R environment to install them:

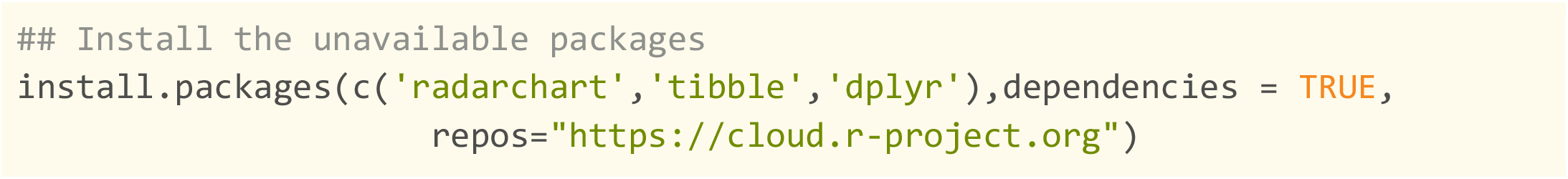

### Installation

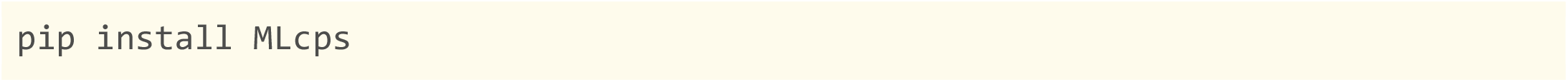

Please refer to https://github.com/FunctionalUrology/MLcps for a detailed tutorial.

## Availability and Implementation

MLcps is available as a Python package at https://pypi.org/project/MLcps/ and the source code is available at https://github.com/FunctionalUrology/MLcps.

MLcps is developed using Python [12] and R [13] programming languages. Pandas [14] is used to store and process the data. Plotly is used to generate the figures. The radarchart [15] package in R was used for surface area calculation of the polar plane.

## Acknowledgment

We gratefully acknowledge the financial support of the Swiss National Science Foundation (SNF Grant 310030_175773 to FCB and KM) and the Wings for Life Spinal Cord Research Foundation (WFL-AT-06/19 to KM). AHG and RMA are supported by R01 DK 077195 and R01 DK127673. MK is supported by the Else Kröner-Fresenius-Stiftung (EKFS 2021_EKeA.33). The assistance provided by Nezhla Aghaei is greatly appreciated for guidance in the mathematical formulation and consultation in the calculation of the planar surface area. The authors acknowledge the financial support from the Federal Ministry of Education and Research of Germany and by the Sächsische Staatsministerium für Wissenschaft Kultur und Tourismus in the program Center of Excellence for AI-research “Center for Scalable Data Analytics and Artificial Intelligence Dresden/Leipzig” (project identification number: ScaDS.AI).

## References

1. Sun, Y.M., A.K.C. Wong, and M.S. Kamel, Classification of Imbalanced Data: A Review. International Journal of Pattern Recognition and Artificial Intelligence, 2009. 23(4): p. 687–719.

2. Huang, J. and C.X. Ling, Using AUC and accuracy in evaluating learning algorithms. Ieee Transactions on Knowledge and Data Engineering, 2005. 17(3): p. 299–310.

3. Huang, J., J.J. Lu, and C.X. Ling, Comparing naive bayes, decision trees, and SVM with AUC and, accuracy. Third Ieee International Conference on Data Mining, Proceedings, 2003: p. 553–556.

4. Provost, F. and P. Domingos, Tree induction for probability-based ranking. Machine Learning, 2003. 52(3): p. 199–215.

5. Racz, A., D. Bajusz, and K. Heberger, Multi-Level Comparison of Machine Learning Classifiers and Their Performance Metrics. Molecules, 2019. 24(15).

6. Branco, P., L. Torgo, and R.P. Ribeiro, A Survey of Predictive Modeling on Im balanced Domains. Acm Computing Surveys, 2016. 49(2).

7. Galar, M., et al., A Review on Ensembles for the Class Imbalance Problem: Bagging-, Boosting-, and Hybrid-Based Approaches. Ieee Transactions on Systems Man and Cybernetics Part C-Applications and Reviews, 2012. 42(4): p. 463–484.

8. Dietrich, S., et al., Drug-perturbation-based stratification of blood cancer. Journal of Clinical Investigation, 2018. 128(1): p. 427–445.

9. Witten, D., et al., Ultra-high throughput sequencing-based small RNA discovery and discrete statistical biomarker analysis in a collection of cervical tumours and matched controls. Bmc Biology, 2010. 8.

10. Colaprico, A., et al., TCGAbiolinks: an R/Bioconductor package for integrative analysis of TCGA data. Nucleic Acids Research, 2016. 44(8).

11. Robinson, M.D., D.J. McCarthy, and G.K. Smyth, edgeR: a Bioconductor package for differential expression analysis of digital gene expression data. Bioinformatics, 2010. 26(1): p. 139–140.

12. Van Rossum, G., & Drake, F. L., Python 3 Reference Manual. 2009.

13. Team, R.C., R: A language and environment for statistical computing. R Foundation for Statistical Computing. 2013.

14. McKinney, Data Structures for Statistical Computing in Python. Proceedings of the 9th Python in Science Conference,, 2010.

15. Porter, D.A.a.S., radarchart: Radar Chart from ‘Chart.js’. R package version 0.3.1. 2016.

