## Supplementary file for "MLcps: Machine Learning Cumulative Performance Score for classification problems"

* Corresponding author:

#### Machine Learning Pipeline

The proposed pipeline (Figure 2) starts by splitting the input datasets into k (3) equal size different bins in a stratified manner, where k-1 bins will be used as training datasets and the remaining bin as a test dataset. Then, it uses the SelectKBest method to select features of a given dataset. With the SelectKBest method, features are selected based on the feature with the highest score. The imbalanced datasets were then sampled with a combined over- and under-sampling method called SMOTETomek [1]. The SMOTETomek method combines the ability of SMOTE to build synthetic data for a minority class with the ability of Tomek Links to remove data from the majority class if they are identified as Tomek links (that is, samples of data from the majority class that are near to the minority class data).

In the next step, the pipeline trains multiple ML algorithms for the given problem. ML algorithms can be broadly categorized into a) Linear Algorithms b) Nonlinear Algorithms and c) Ensemble algorithms. Here, we have selected at least 1 classification algorithm from each of these categories. In total 7 (Table S1) different classification algorithms were trained for each dataset. In addition, a Dummy classifier that makes predictions at random was used as a baseline to compare with other models.

*Table S1: ML algorithms used in this study*

| **Classification Algorithm** | **Abbreviation** |
|---|---|
| Logistic Regression | LR |
| Linear Discriminant Analysis | LDA |
| Support Vector Machine | SVM |
| Gaussian Process Classifier | GP |
| Random Forest Classifier | RF |
| Bagging Classifier | BC |
| Extra Trees Classifier | ETC |

Then, the performance of the trained ML models was evaluated using the *k-fold cross-validation (CV) (where k=3)* method which divides the whole dataset into *k* non-overlapping subsets of equal size. For each fold, ($k - 1$) subsets are used as a training dataset for the model, and the remaining subset as a test dataset to evaluate the model performance. In this way, it trains k different models and the final performance of the model is estimated by taking the average of the evaluation metrics from each iteration. Since a single run of the k-fold cross-validation method may result in a noisy estimate of model performance, we repeated (n) k-fold CV 10 times.

Additionally, larger TCGA-BRCA datasets allow us to create an independent test set (30% of the datasets). Therefore, we have measured the performance of the trained model on unseen data as well for TCGA-BRCA datasets. Similar to ML algorithms, several different performance metrics that *are Accuracy, Balanced Accuracy, Precision, Recall, Average Precision, and ROC-AUC score,* were used to evaluate the performance of ML models. The complete pipeline was developed on top of the sci-kit-learn library [2] and imblearn [3] from Python . Pandas package was used to store and process the data.

#### Quick Start for MLcps in python

```
from MLcps import getCPS

#calculate Machine Learning cumulative performance score
cps=getCPS.calculate(object)
```

where:

***object***: A pandas dataframe where rows are different metrics scores and columns are different ML models. Or a GridSearchCV object.

***cps***: A pandas dataframe with model's name and corresponding MLcps. Or a GridSearchCV object.

#### Weighted MLcps

```
#read input data (a dataframe) or load an example data
metrics=getCPS.load_metrics()

#define weights
weights={"Accuracy":0.75,"F1": 1.25}

#calculate Machine Learning cumulative performance score
cpsScore=getCPS.calculate(metrics,weights)

########################################################
#load GridSearch object or load it from package
```

```
gsObj=getCPS.load_GridSearch_Object()

#define weights
weights={"accuracy":0.75,"f1": 1.25}

#calculate Machine Learning cumulative performance score
gsObj_updated=getCPS.calculate(gsObj,weights)
```

Please refer to https://github.com/FunctionalUrology/MLcps for a detailed tutorial.

#### Python (ML pipeline) Session Information

```
-----
imblearn          0.9.1
ipykernel         6.13.0
joblib            1.1.0
matplotlib        3.5.2
numpy             1.22.3
pandas            1.4.2
plotly            5.8.0
session_info      1.0.0
sklearn           1.1.0
-----
IPython           8.3.0
jupyter_client    7.3.1
jupyter_core      4.10.0
-----
Python 3.10.4 | packaged by conda-forge | (main, Mar 24 2022, 17:39:04) [GCC 10.3.0]
Linux-3.10.0-1160.62.1.el7.x86_64-x86_64-with-glibc2.17
-----
Session information updated at 2022-06-08 17:31
```

#### Packages and other dependencies used in MLcps

##### Python

```
-----
MLcps             0.0.5
numpy             1.22.3
pandas            1.4.2
pkg_resources     NA
plotly            5.8.0
rpy2              3.5.1
session_info      1.0.0
```

-----
IPython 7.33.0
jupyter_client 7.3.1
jupyter_core 4.10.0
-----
Python 3.10.4 | packaged by conda-forge | (main, Mar 24 2022, 17:45:10) [Clang 12.0.1 ]
macOS-11.2.3-x86_64-i386-64bit
-----
Session information updated at 2022-06-08 10:07

## R

| Package | Version | Reference |
|---|---|---|
| **tidyselect** | 1.1.2 | Lionel Henry and Hadley Wickham (2022). tidyselect: Select from a Set of Strings. R package version 1.1.2. https://CRAN.R-project.org/package=tidyselect |
| **purrr** | 0.3.4 | Lionel Henry and Hadley Wickham (2020). purrr: Functional Programming Tools. R package version 0.3.4. https://CRAN.R-project.org/package=purrr |
| **haven** | 2.4.1 | Hadley Wickham and Evan Miller (2021). haven: Import and Export 'SPSS', 'Stata' and 'SAS' Files. R package version 2.4.1. https://CRAN.R-project.org/package=haven |
| **carData** | 3.0.4 | John Fox, Sanford Weisberg and Brad Price (2020). carData: Companion to Applied Regression Data Sets. R package version 3.0-4. https://CRAN.R-project.org/package=carData |
| **colorspace** | 2.0.2 | Not available |
| **vctrs** | 0.4.1 | Hadley Wickham, Lionel Henry and Davis Vaughan (2022). vctrs: Vector Helpers. R package version 0.4.1. https://CRAN.R-project.org/package=vctrs |
| **generics** | 0.1.2 | Hadley Wickham, Max Kuhn and Davis Vaughan (2022). generics: Common S3 Generics not Provided by Base R Methods Related to |
| **htmltools** | 0.5.1.1 | Joe Cheng, Carson Sievert, Winston Chang, Yihui Xie and Jeff Allen (2021). htmltools: Tools for HTML. R package version 0.5.1.1. https://CRAN.R-project.org/package=htmltools |
| **utf8** | 1.2.1 | Patrick O. Perry (2021). utf8: Unicode Text Processing. R package version 1.2.1. https://CRAN.R-project.org/package=utf8 |
| **rlang** | 1.0.2 | Lionel Henry and Hadley Wickham (2022). rlang: Functions for Base Types and Core R and 'Tidyverse' Features. R package version 1.0.2. https://CRAN.R-project.org/package=rlang |
| **pillar** | 1.7.0 | Kirill Müller and Hadley Wickham (2022). pillar: Coloured Formatting for Columns. R package version 1.7.0. https://CRAN.R-project.org/package=pillar |

| | | |
|---|---|---|
| **foreign** | 0.8.81 | R Core Team (2020). foreign: Read Data Stored by 'Minitab', 'S', 'SAS', 'SPSS', 'Stata', |
| **glue** | 1.6.2 | Jim Hester and Jennifer Bryan (2022). glue: Interpreted String Literals. R package version 1.6.2. https://CRAN.R-project.org/package=glue |
| **withr** | 2.4.2 | Jim Hester, Kirill Müller, Kevin Ushey, Hadley Wickham and Winston Chang (2021). withr: Run Code 'With' Temporarily Modified Global State. R package version 2.4.2. https://CRAN.R-project.org/package=withr |
| **DBI** | 1.1.1 | R Special Interest Group on Databases (R-SIG-DB), Hadley Wickham and Kirill Müller (2021). DBI: R Database Interface. R package version 1.1.1. https://CRAN.R-project.org/package=DBI |
| **RColorBrewer** | 1.1.2 | Erich Neuwirth (2014). RColorBrewer: ColorBrewer Palettes. R package version 1.1-2. https://CRAN.R-project.org/package=RColorBrewer |
| **readxl** | 1.3.1 | Hadley Wickham and Jennifer Bryan (2019). readxl: Read Excel Files. R package version 1.3.1. https://CRAN.R-project.org/package=readxl |
| **lifecycle** | 1.0.1 | Lionel Henry and Hadley Wickham (2021). lifecycle: Manage the Life Cycle of your Package Functions. R package version 1.0.1. https://CRAN.R-project.org/package=lifecycle |
| **plyr** | 1.8.6 | Hadley Wickham (2011). The Split-Apply-Combine Strategy for Data Analysis. Journal of Statistical Software, 40(1), 1-29. URL http://www.jstatsoft.org/v40/i01/. |
| **cellranger** | 1.1.0 | Jennifer Bryan (2016). cellranger: Translate Spreadsheet Cell Ranges to Rows and Columns. R package version 1.1.0. https://CRAN.R-project.org/package=cellranger |
| **munsell** | 0.5.0 | Charlotte Wickham (2018). munsell: Utilities for Using Munsell Colours. R package version 0.5.0. https://CRAN.R-project.org/package=munsell |
| **ggsignif** | 0.6.2 | Ahlmann-Eltze, C., & Patil, I. (2021). ggsignif: R Package for Displaying Significance Brackets for 'ggplot2'. PsyArxiv. doi:10.31234/osf.io/7awm6 |
| **gtable** | 0.3.0 | Hadley Wickham and Thomas Lin Pedersen (2019). gtable: Arrange 'Grobs' in Tables. R package version 0.3.0. https://CRAN.R-project.org/package=gtable |
| **zip** | 2.2.0 | Gábor Csárdi, Kuba Podgórski and Rich Geldreich (2021). zip: Cross-Platform 'zip' Compression. R package version 2.2.0. https://CRAN.R-project.org/package=zip |
| **htmlwidgets** | 1.5.3 | Ramnath Vaidyanathan, Yihui Xie, JJ Allaire, Joe Cheng, Carson Sievert and Kenton Russell (2020). htmlwidgets: HTML Widgets for R. R package version 1.5.3. https://CRAN.R-project.org/package=htmlwidgets |
| **forcats** | 0.5.1 | Hadley Wickham (2021). forcats: Tools for Working with Categorical Variables (Factors). R package version 0.5.1. https://CRAN.R-project.org/package=forcats |
| **rio** | 0.5.27 | Chung-hong Chan, Geoffrey CH Chan, Thomas J. Leeper, and Jason Becker (2021). rio: A Swiss-army knife for data file I/O. R package version 0.5.27. |
| **extrafont** | 0.17 | Winston Chang, (2014). extrafont: Tools for using fonts. R package version 0.17. https://CRAN.R-project.org/package=extrafont |
| **curl** | 4.3.2 | Jeroen Ooms (2021). curl: A Modern and Flexible Web Client for R. R package version 4.3.2. https://CRAN.R-project.org/package=curl |

| | | |
|---|---|---|
| **fansi** | 0.5.0 | Brodie Gaslam (2021). fansi: ANSI Control Sequence Aware String Functions. R package version 0.5.0. https://CRAN.R-project.org/package=fansi |
| **Rttf2pt1** | 1.3.8 | Winston Chang, Andrew Weeks, Frank M. Siegert, Mark Heath, Thomas Henlick, Sergey Babkin, Turgut Uyar, Rihardas Hepas, Szalay Tamas, Johan Vromans, Petr Titera, Lei Wang, Chen Xiangyang, Zvezdan Petkovic, Rigel and I. Lee Hetherington (2020). Rttf2pt1: 'ttf2pt1' Program. R package version 1.3.8. https://CRAN.R-project.org/package=Rttf2pt1 |
| **broom** | 0.7.12 | David Robinson, Alex Hayes and Simon Couch (2022). broom: Convert Statistical Objects into Tidy Tibbles. R package version 0.7.12. https://CRAN.R-project.org/package=broom |
| **Rcpp** | 1.0.7 | Dirk Eddelbuettel and Romain Francois (2011). Rcpp: Seamless R and C++ Integration. Journal of Statistical Software, 40(8), 1-18. URL https://www.jstatsoft.org/v40/i08/. |
| **KernSmooth** | 2.23.18 | Matt Wand (2020). KernSmooth: Functions for Kernel Smoothing Supporting Wand & Jones (1995). R package version 2.23-18. https://CRAN.R-project.org/package=KernSmooth |
| **backports** | 1.2.1 | Michel Lang and R Core Team (2020). backports: Reimplementations of Functions Introduced Since R-3.0.0. R package version 1.2.1. https://CRAN.R-project.org/package=backports |
| **abind** | 1.4.5 | Tony Plate and Richard Heiberger (2016). abind: Combine Multidimensional Arrays. R package version 1.4-5. https://CRAN.R-project.org/package=abind |
| **proj4** | 1.0.10.1 | Simon Urbanek (2021). proj4: A simple interface to the PROJ.4 cartographic projections |
| **hms** | 1.1.0 | Kirill Müller (2021). hms: Pretty Time of Day. R package version 1.1.0. https://CRAN.R-project.org/package=hms |
| **digest** | 0.6.27 | Dirk Eddelbuettel with contributions by Antoine Lucas, Jarek Tuszynski, Henrik Bengtsson, Simon Urbanek, Mario Frasca, Bryan Lewis, Murray Stokely, Hannes Muehleisen, Duncan Murdoch, Jim Hester, Wush Wu, Qiang Kou, Thierry Onkelinx, Michel Lang, Viliam Simko, Kurt Hornik, Radford Neal, Kendon Bell, Matthew de Queljoe, Ion Suruceanu, Bill Denney, Dirk Schumacher and and Winston Chang. (2020). digest: Create Compact Hash Digests of R Objects. R package version 0.6.27. https://CRAN.R-project.org/package=digest |
| **openxlsx** | 4.2.4 | Philipp Schauberger and Alexander Walker (2021). openxlsx: Read, Write and Edit xlsx Files. R package version 4.2.4. https://CRAN.R-project.org/package=openxlsx |
| **stringi** | 1.7.2 | Not available |
| **rstatix** | 0.7.0 | Alboukadel Kassambara (2021). rstatix: Pipe-Friendly Framework for Basic Statistical Tests. R package version 0.7.0. https://CRAN.R-project.org/package=rstatix |
| **ash** | 1.0.15 | S original by David W. Scott R port by Albrecht Gebhardt adopted to recent S-PLUS by Stephen Kaluzny <> (2015). ash: David Scott's ASH Routines. R package version 1.0-15. https://CRAN.R-project.org/package=ash |
| **grid** | 4.0.5 | R Core Team (2021). R: A language and environment for statistical computing. R Foundation for Statistical Computing, Vienna, Austria. URL https://www.R-project.org/. |

| | | |
|---|---|---|
| **cli** | 3.2.0 | Gábor Csárdi (2022). cli: Helpers for Developing Command Line Interfaces. R package version 3.2.0. https://CRAN.R-project.org/package=cli |
| **tools** | 4.0.5 | R Core Team (2021). R: A language and environment for statistical computing. R Foundation for Statistical Computing, Vienna, Austria. URL https://www.R-project.org/. |
| **magrittr** | 2.0.1 | Stefan Milton Bache and Hadley Wickham (2020). magrittr: A Forward-Pipe Operator for R. R package version 2.0.1. https://CRAN.R-project.org/package=magrittr |
| **maps** | 3.3.0 | Original S code by Richard A. Becker, Allan R. Wilks. R version by Ray Brownrigg. Enhancements by Thomas P Minka and Alex Deckmyn. (2018). maps: Draw Geographical Maps. R package version 3.3.0. https://CRAN.R-project.org/package=maps |
| **crayon** | 1.4.1 | Gábor Csárdi (2021). crayon: Colored Terminal Output. R package version 1.4.1. https://CRAN.R-project.org/package=crayon |
| **extrafontdb** | 1.0 | Winston Chang (2012). extrafontdb: Package for holding the database for the extrafont package. R package version 1.0. https://CRAN.R-project.org/package=extrafontdb |
| **car** | 3.0.11 | John Fox and Sanford Weisberg (2019). An [4] Companion to Applied Regression, Third Edition. Thousand Oaks CA: Sage. URL: https://socialsciences.mcmaster.ca/jfox/Books/Companion/ |
| **pkgconfig** | 2.0.3 | Gábor Csárdi (2019). pkgconfig: Private Configuration for 'R' Packages. R package version 2.0.3. https://CRAN.R-project.org/package=pkgconfig |
| **ellipsis** | 0.3.2 | Hadley Wickham (2021). ellipsis: Tools for Working with .... R package version 0.3.2. https://CRAN.R-project.org/package=ellipsis |
| **MASS** | 7.3.53.1 | Venables, W. N. & Ripley, B. D. (2002) Modern Applied Statistics with S. Fourth Edition. Springer, New York. ISBN 0-387-95457-0 |
| **data.table** | 1.14.0 | Matt Dowle and Arun Srinivasan (2021). data.table: Extension of \`data.frame\`. R package version 1.14.0. https://CRAN.R-project.org/package=data.table |
| **assertthat** | 0.2.1 | Hadley Wickham (2019). assertthat: Easy Pre and Post Assertions. R package version 0.2.1. https://CRAN.R-project.org/package=assertthat |
| **R6** | 2.5.0 | Winston Chang (2020). R6: Encapsulated Classes with Reference Semantics. R package version 2.5.0. https://CRAN.R-project.org/package=R6 |
| **compiler** | 4.0.5 | R Core Team (2021). R: A language and environment for statistical computing. R Foundation for Statistical Computing, Vienna, Austria. URL https://www.R-project.org/. |
| **dplyr** | 1.0.9 | Hadley Wickham, Romain François, Lionel Henry and Kirill Müller (2022). dplyr: A Grammar of Data Manipulation. R package version 1.0.9. https://CRAN.R-project.org/package=dplyr |
| **tidyr** | 1.2.0 | Hadley Wickham and Maximilian Girlich (2022). tidyr: Tidy Messy Data. R package version 1.2.0. https://CRAN.R-project.org/package=tidyr |
| **tibble** | 3.1.7 | Kirill Müller and Hadley Wickham (2022). tibble: Simple Data Frames. R package version 3.1.7. https://CRAN.R-project.org/package=tibble |

| | | |
|---|---|---|
| **ggpubr** | 0.4.0 | Alboukadel Kassambara (2020). ggpubr: 'ggplot2' Based Publication Ready Plots. R package version 0.4.0. https://CRAN.R-project.org/package=ggpubr |
| **ggalt** | 0.4.0 | Bob Rudis, Ben Bolker and Jan Schulz (2017). ggalt: Extra Coordinate Systems, 'Geoms', Statistical Transformations, |
| **ggplot2** | 3.3.5 | H. Wickham. ggplot2: Elegant Graphics for Data Analysis. Springer-Verlag New York, 2016. |
| **ggridges** | 0.5.3 | Claus O. Wilke (2021). ggridges: Ridgeline Plots in 'ggplot2'. R package version 0.5.3. https://CRAN.R-project.org/package=ggridges |
| **stringr** | 1.4.0 | Hadley Wickham (2019). stringr: Simple, Consistent Wrappers for Common String Operations. R package version 1.4.0. https://CRAN.R-project.org/package=stringr |
| **scales** | 1.2.0 | Hadley Wickham and Dana Seidel (2022). scales: Scale Functions for Visualization. R package version 1.2.0. https://CRAN.R-project.org/package=scales |
| **radarchart** | 0.3.1 | Doug Ashton and Shane Porter (2016). radarchart: Radar Chart from 'Chart.js'. R package version 0.3.1. https://CRAN.R-project.org/package=radarchart |

### References:


1. G. Batista, B.B., M. Monard, *Balancing Training Data for Automated Annotation of Keywords: a Case Study.* WOB, 2003.
2. Pedregosa, F., et al., *Scikit-learn: Machine Learning in Python.* Journal of Machine Learning Research, 2011. **12**: p. 2825-2830.
3. Lemaitre, G., F. Nogueira, and C.K. Aridas, *Imbalanced-learn: A Python Toolbox to Tackle the Curse of Imbalanced Datasets in Machine Learning.* Journal of Machine Learning Research, 2017. **18**.
